# Serum metabolomics reveals signatures associated with physical resilience trajectories from middle to older age

**DOI:** 10.64898/2026.09.14.751497

**Authors:** Jeong In Seo, Toon A.W. Scheurink, Crystal X. Wang, Kine Eide Kvitne, Wilhan D. Gonçalves Nunes, Jasmine Zemlin, Jaclyn Bergstrom, Pieter C. Dorrestein, Ipsita Mohanty, Anthony J.A. Molina

## Abstract

Lifecourse physical resilience is defined by the ability to maintain abilities across multiple domains of physical performance. While the importance of physical resilience in functional independence and mobility disability is clear, studies investigating metabolomic signatures of physical resilience are lacking. Here, we performed untargeted metabolomics on serum samples from a community-based cohort of 237 individuals followed over 28 years, and applied spectral data mining tools to map identified metabolites to health phenotypes from public repositories. We identified metabolites across multiple chemical classes, including acylcarnitines, glutamine conjugates, and phosphocholines, that were differentially associated with physical resilience status. Notably, medium-chain acylcarnitines negatively associated with physical resilience were more frequently observed in disease phenotypes than in healthy individuals. Kynurenine, a tryptophan metabolite linked to age-related functional decline, increased more steeply with age in individuals with low physical resilience. We also found that metabolites of the antihypertensive drug verapamil were associated with physical resilience in a metabolism-dependent manner, differing between oxidative and glucuronidated forms. Together, these metabolic signatures offer a resource for identifying biochemical pathways and biomarkers relevant to physical resilience for healthy aging.

## Introduction

Changes in physical ability with advancing age underlie the progression of mobility disability and loss of functional independence. As the population of older adults grows globally due to increased life expectancy, the burden of functional decline and frailty among continues to rise^1,2^. Importantly, aging trajectories vary substantially between individuals, suggesting that there are opportunities to modify the trajectory of progressive physical function decline^3^. While chronological age affects average declines in physical function, some individuals maintain robust physical function well into late life, whereas others experience expected or accelerated physical deterioration. Understanding the molecular basis of this variability may help identify biological pathways involved in preserving physical function during aging.

Physical resilience, defined as the capacity to maintain or recover function in response to chronological aging or health-related stressors, declines with advancing age and is therefore of particular relevance to older adult populations^4,5^. In prior cohort studies, higher levels of physical resilience have been consistently associated with reduced risk of adverse health outcomes, including mortality, falls, and hospitalization, with evidence suggesting wide ranging benefits as resilience levels increase^6^. Notably, this protective association extends even to individuals with lifespan-limiting genetic risk factors, highlighting the potential importance of physical resilience as a determinant of healthy aging^7^.

Untargeted metabolomics provides a means to systematically characterize molecular processes associated with aging-related phenotypes because the metabolome reflects the combined influence of genetic, environmental, lifestyle, and disease-related factors^8^. However, metabolomic studies directly examining physical resilience over time remain scarce. One related study characterized the metabolomic signatures of physical frailty, but relied on a targeted approach restricted to predefined lipid classes, finding several ceramide species elevated in physically frail compared to robust older adults^9^. Such targeted approaches, however, are inherently restricted to pre-selected biochemical classes and may overlook novel pathways relevant to physical resilience.

To identify metabolite signatures associated with physical resilience, we performed Liquid Chromatography-Tandem Mass Spectrometry (LC-MS/MS)-based untargeted serum metabolomics in the Rancho Bernardo Study (RBS), a community-based cohort^10^ in which metabolomic correlates of cognitive resilience have previously been characterized^11^. We further leveraged data mining tools within the Global Natural Product Social Molecular Networking 2 (GNPS2) ecosystem to assess the occurrence of resilience-associated metabolites across publicly available mass spectrometry datasets, providing broader biological context for identified features^12,13^. Together, these analyses identify metabolite signatures associated with physical resilience and provide a foundation for future studies aimed at elucidating mechanisms of healthy aging.

## Results

### Cohort characterization and analysis pipeline

Serum samples for metabolomics analysis were obtained from 237 participants in the prospective longitudinal Rancho Bernardo Study (RBS) of Healthy Aging, which was previously used for metabolomics profiling of cognitive resilience^10,11^. RBS has extensive metadata, including clinical, physical, and demographic outcomes (**Figure 1a**). For this study, we utilized the physical resilience scores, calculated based on a participant’s performance on the Grip Strength test and the Timed Chair stand^14^, over 28 years in a subset of 2,630 participants (*see methods for calculation of resilience scores*). Detailed information on the calculation of the physical resilience scores can be found in a previous publication^15^, and they are handled as per^11^.

**Figure 1.**
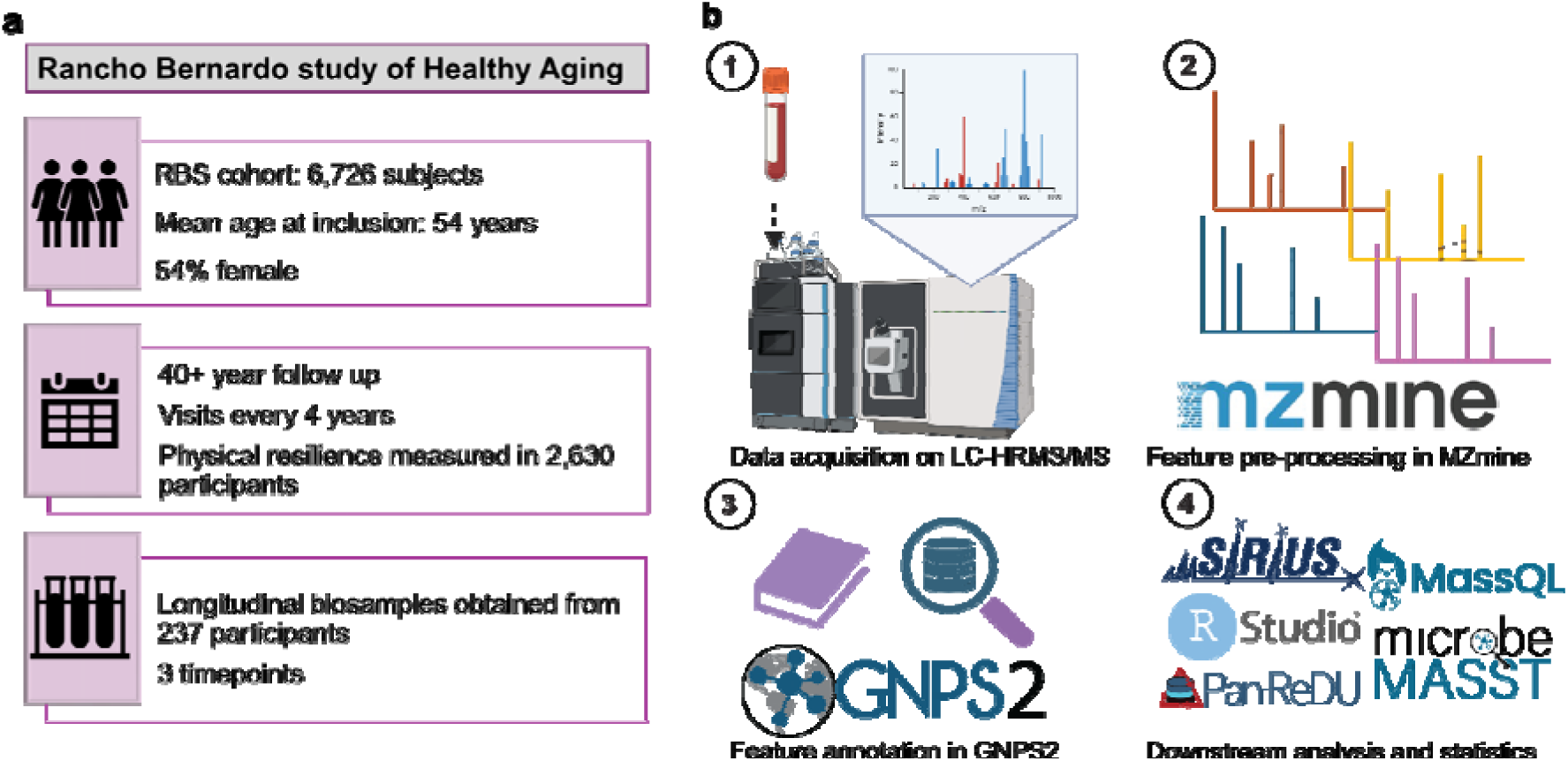
Study outline and workflow. **a)** The Rancho Bernardo study description. **b)** Sample processing and feature detection methods employed, as well as the utilized downstream analysis tools. Icons were obtained from Powerpoint and Biorender, and logos from their respective websites.

We utilized serum samples collected at 3 time points within RBS for 237 participants. The serum samples were analyzed using untargeted LC-MS/MS-based metabolomics. The complete workflow for the metabolomics analysis is shown in **Figure 1b**. To investigate the molecular underpinnings of physical resilience, we selected the first visit in this subset for each participant. In this subset, shown in **Supplementary Table 1**, participants had a mean age of 64.3 (SD: 7.2), had a mean physical resilience score of -0.003 (SD: 0.051), and 152 participants (64%) were female. Participants were dichotomized using a median split into high- and low-resilience groups (physical resilience score: median 0.029 in the high resilience group vs median -0.032 in the low resilience group). The high resilience group contained a higher proportion of females than the low resilience group (76% vs 52%, *p* < 0.001), and had lower BMI (24.82, SD 3.35 vs 25.86, SD 3.53; *p* = 0.012). The metabolite associations were robust to both. All features that differed significantly between resilience groups showed the same direction of effect within females and within males, effect sizes were closely correlated between sexes (Pearson *r* = 0.90), and all remained significant when the comparison was stratified by sex (**Supplementary Table 2**). The BMI difference was present only when both sexes were analysed together and was absent within each sex (females *p* = 0.4; males *p* = 0.2), consistent with it reflecting the sex distribution. Neither variable was therefore included as a covariate.

To identify metabolites associated with physical resilience, we performed sparse partial least squares regression (sPLS-R) on the untargeted LC-MS/MS data. Optimal sparsity parameters were determined using a ranked scree-style plot and cross-validation, resulting in the selection of 2,810 features and 1 component, indicating that a single multivariate dimension best captured the association between metabolomic features and physical resilience. The model explained 10% of the total variance in physical resilience (p < 0.001) (**Figure 2a)**. Observed R^2^ using 10-fold cross-validation (cv) was 0.076 (root mean square error (RMSE) = 0.050). The selected features showed good selection stability, with a median stability score of 0.9 (range: 0.1-1.0), with 82% of features selected in at least 7 of the 10 cross-validation folds (**Supplementary Figure S1**). From a total of 22,736 features, our model selected 2,810 (12.4%) features (**Figure 2b**). The 2,810 features, their weights, and stability can be found in **Supplementary Table 3**.

**Figure 2.**
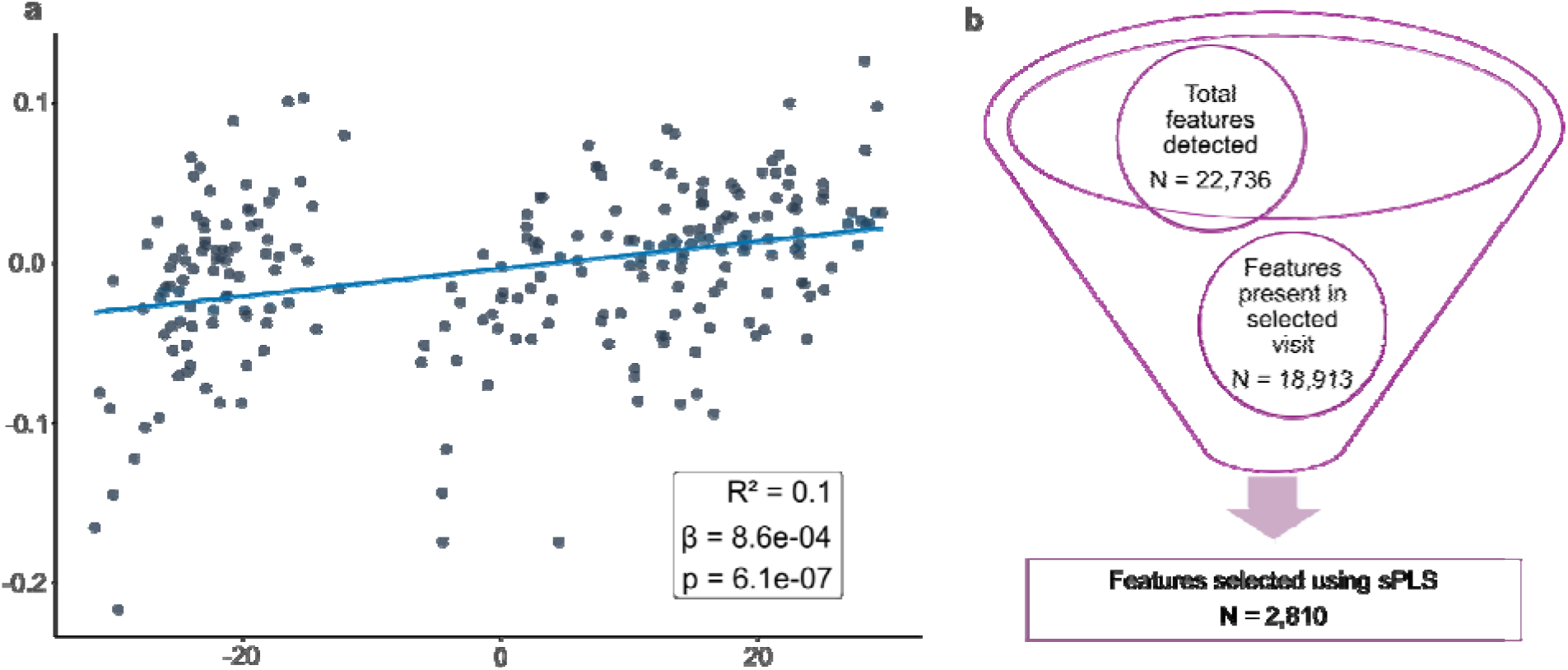
Model performance and selection of resilience-associated features. **a)** A linear regression of the first component of the sPLS model against the physical resilience scores. **b)** Selection of the 2,810 features associated with physical resilience. Icons were obtained from Powerpoint.

### Molecular Networking Uncovers Chemical Classes Associated with Physical Resilience

To obtain an overview of the compounds associated with physical resilience, we mapped them as molecular networks^16^, in which compounds with similar MS/MS spectral patterns are connected to one another, such that compounds belonging to the same chemical class are likely to fall within the same (sub)network^17^. **Figure 3** shows a composite of the key molecular networks derived from this dataset, where node size corresponds to the weight from the sPLS model and node color represents the association with physical resilience (blue indicating a positive association and yellow indicating a negative association). As in our previous study of the serum metabolomic signatures associated with cognitive resilience^11^, we again observed that the largest network consisted of acylcarnitines of varying chain lengths, with medium-chain (C6–C12) acylcarnitines^18^ being the most prevalent type negatively associated with physical resilience. For other compound classes negatively associated with physical resilience selected by the sPLS-R model, we observed MS/MS spectral matches to tryptophan metabolites from both the kynurenine and indole pathways, such as xanthurenic acid, 4-hydroxy-2-quinolinecarboxylic acid (also known as kynurenic acid), and indole-3-acetic acid, as well as glutamine conjugates, most of which were conjugated with acyl chains of varying chain length and saturation. In contrast, we found that the sub-network of phosphocholines was positively associated with physical resilience. Notably, several classes of anti-hypertensive drugs were associated negatively with physical resilience. To investigate if these medications change the association of other molecular classes with resilience, we re-ran the model in participants where no anti-hypertensive drugs were detected (n = 202). Model performance was highly similar (observed R^2^ = 0.13, **Supplementary Figure S2a**) and the selected metabolites by this model and the original one were strongly correlated (r = 0.918, **Supplementary Figure S2b**). For features without library annotation, due to the limited spectral library coverage currently available on GNPS2, SIRIUS^19^, an *in silico* tool that classifies each metabolite into a specific NPC (natural product compound) superclass, was employed to examine the distribution of unannotated features across NPC superclasses (**Figure 3b**). Among these, glycerophospholipids, the superclass containing phosphocholines, showed a predominantly positive association with physical resilience, consistent with the positive association observed for the phosphocholine molecular network described above.

**Figure 3.**
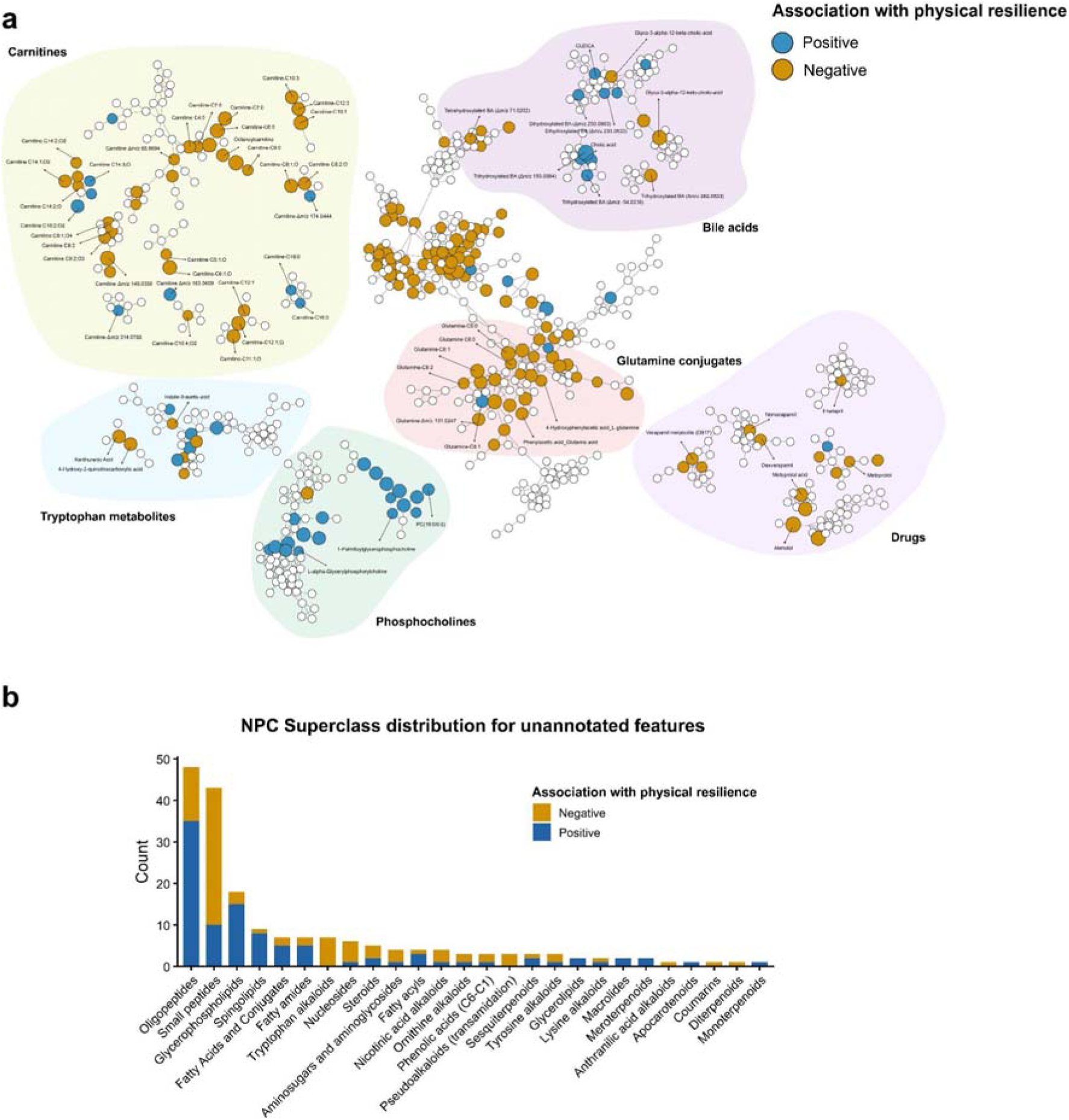
Metabolite overview of features associated with physical resilience. (a) Molecular network of annotated features associated with physical resilience on GNPS2. (b) Distribution of different chemical classes among unannotated features, determined by SIRIUS 6.3.2.

Furthermore, the features associated with physical resilience overlapped only partially with those we previously reported for cognitive resilience^11^, in both the negatively and positively associated directions, suggesting that key features across these two domains of resilience may be driven by distinct metabolic signatures **(Supplementary Figure S3a and S3b).**

### Acylcarnitines, Glutamine Conjugates, and Phosphocholines Associated with Physical Resilience Show Distinct Disease Distribution in Public Repositories

To characterize the individual features driving the class-specific associations identified in the molecular networks (**Figure 3a**), we examined the metabolites that differed significantly between high- and low-physical-resilience groups within each class. Participants in the RBS cohort were divided into these groups based on quartiles of physical resilience scores, with the lower two quartiles assigned to the low-resilience group and the upper two quartiles to the high-resilience group; significant differences were assessed using the Wilcoxon rank-sum test with Benjamini–Hochberg (BH) correction for multiple comparisons. Among the acylcarnitines, a broad set of species was depleted in the high-resilience group (**Figure 4a**), including the long-chain acylcarnitine C14:2 (Wilcoxon rank-sum test, BH-adjusted *p* = 0.031; Cliff’s δ = -0.15 [95% CI: -0.25, -0.06]), the medium-chain acylcarnitine C6:1;O2 (Wilcoxon rank-sum test, BH-adjusted *p* = 0.027; Cliff’s δ = -0.21 [95% CI: -0.32, -0.11]), and an unannotated feature with a delta mass of 270.1678 (Wilcoxon rank-sum test, BH-adjusted *p* = 0.031; Cliff’s delta, δ = -0.24 [95% CI: -0.37, -0.09]). The glutamine conjugates followed the same direction, with every significant species, including glutamine C11:1 (Wilcoxon rank-sum test, BH-adjusted *p* = 0.009; Cliff’s delta, δ = -0.16 [95% CI: -0.25, -0.07]) and two glutamine C8:2 isomers at distinct retention times C8:2 (RT -2.58 min; Wilcoxon rank-sum test, BH-adjusted *p* = 0.027; Cliff’s delta, δ = -0.15 [95% CI: -0.26, -0.05]) and C8:2 (RT -3.59 min; Wilcoxon rank-sum test, BH-adjusted *p* = 0.04; Cliff’s delta, δ = -0.16 [95% CI: -0.28, -0.03]), depleted in the high-resilience group (**Figure 4b**). The phosphocholines, by contrast, were enriched in high-resilience individuals (**Figure 4c**), including putative PC(20:2/0:0) (Wilcoxon rank-sum test, BH-adjusted *p* = 0.018; Cliff’s delta, δ = 0.27 [95% CI: 0.13, 0.39]) and putative SM(d18:1/12:0) (Wilcoxon rank-sum test, BH-adjusted *p* = 0.039; Cliff’s delta, δ = 0.25 [95% CI: 0.10, 0.38]), along with several other sphingomyelin (SM) species.

**Figure 4.**
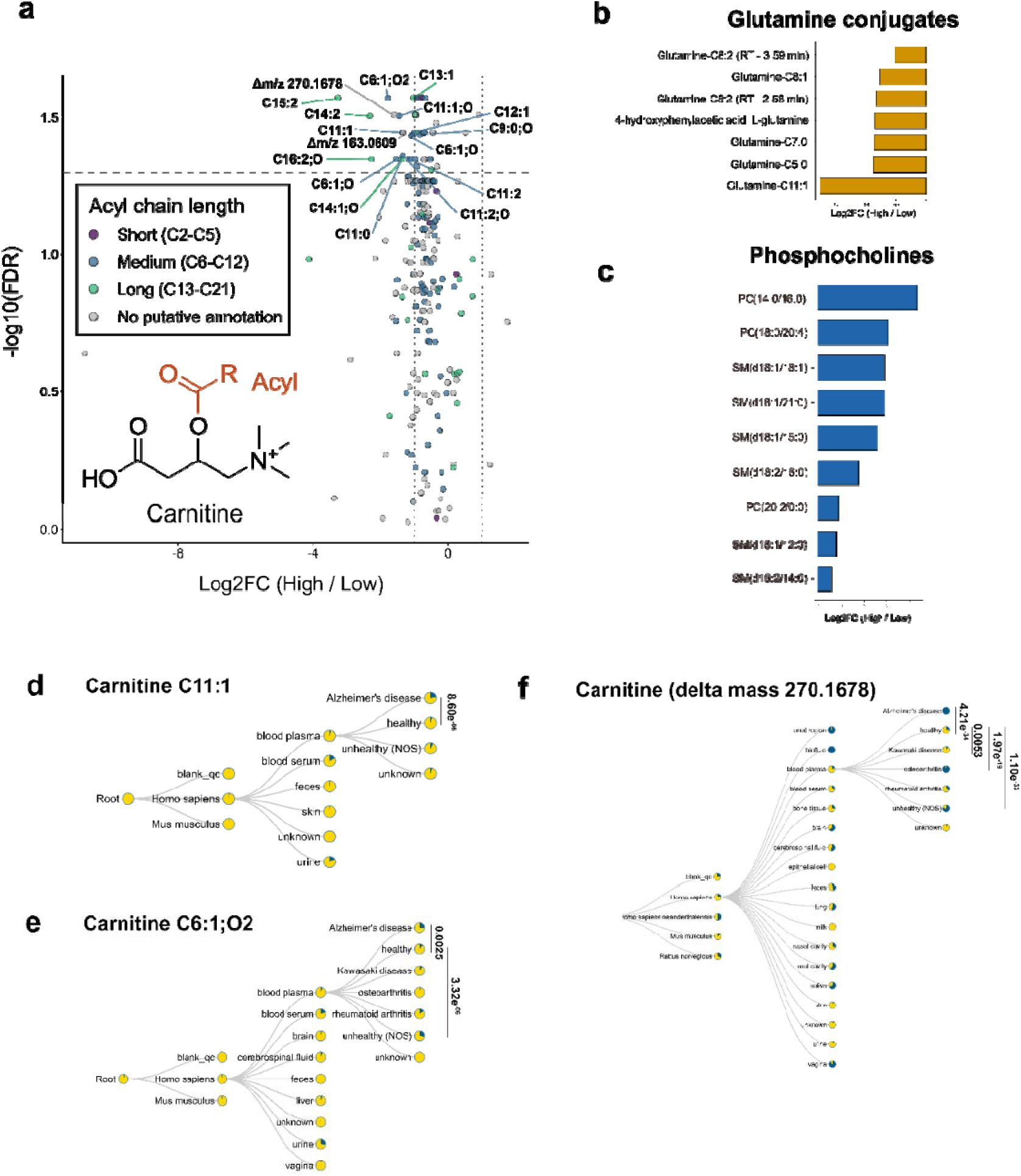
Untargeted metabolomics identifies acylcarnitines, glutamine conjugates, and phosphocholines associated with physical resilience and reveals their disease distributions in humans. (a) A volcano plot showing acylcarnitines with statistically significant differences between groups. (b-c) Bar plots showing metabolites significantly enriched or depleted in individuals with high versus low physical resilience, including (b) glutamine conjugates, and (c) phosphocholines (Benjamini–Hochberg adjusted *P* < 0.05 and log2FC > 0.5 for glutamine conjugates and phosphocholines). Values represent log2 fold changes (high-resilience group/low-resilience group). (d) C11:1 carnitine, (e) C6:1;O2 carnitine, and (f) carnitine (delta mass 270.1678) were searched across publicly available datasets using tissueMASST. Detection rates were compared across each disease, with statistically significant associations determined using Fisher’s exact test. Metabolites were putatively annotated by spectral library matching in GNPS2. Individuals were classified into high- and low-physical-resilience groups based on quartiles of physical resilience scores within the complete RBS cohort, with the lower two quartiles assigned to the low-resilience group and the upper two quartiles assigned to the high-resilience group.

We next asked whether the resilience-associated features had documented disease links in public data, querying representative features against publicly available metabolomics datasets with tissueMASST^20^. Although not all features were well represented in these datasets, several acylcarnitines showed disease-specific occurrence in humans. Carnitine C11:1 (Fisher’s exact test, *p* = 8.60e-06), carnitine C6:1;O2 (Fisher’s exact test, *p* = 0.0025), and the unannotated feature at delta mass 270.1678 (Fisher’s exact test, *p* = 4.21e-34) were detected in human plasma, where their detection rates were significantly higher in individuals with Alzheimer’s disease than in healthy individuals (**Figure 4d-f**). Carnitine C6:1;O2 was detected more often in unhealthy samples, and the delta mass 270.1678 feature was additionally elevated in osteoarthritis, rheumatoid arthritis, Kawasaki disease, and unhealthy individuals (Fisher’s exact test, all *p*<0.05) (**Figure 4e and f**), suggesting that these medium-chain acylcarnitine features recur across several inflammatory and neurodegenerative conditions. Given their depletion in high-resilience individuals, this pattern points to medium-chain acylcarnitine accumulation as a feature shared across physical resilience in aging and a range of disease states. Furthermore, as certain acylcarnitines and phosphocholines were negatively and positively cross-sectionally associated with physical resilience, respectively, we investigated how they changed over time. We selected 3 acylcarnitines and 2 phosphocholines that were found in < 70% of samples and applied a linear mixed model to quantify their abundance with age, which can be found in **Supplementary Table 4**, and the distribution of 2 carnitines across age and visits are shown in **Supplementary Figure S4.**

### Longitudinal Analysis Reveals a Divergent Age Trajectory of Kynurenine with Physical Resilience

The molecular network contained a subnetwork of tryptophan metabolites spanning both the kynurenine and indole pathways, most of which were negatively associated with physical resilience (**Figure 3a**). Tryptophan metabolites have been implicated in age-related decline in physical function, but existing evidence comes largely from animal models or from measurements at a single time point^21–23^. Hence, we further examined whether tryptophan metabolites differed between physical resilience groups in how they changed with advancing age.

Of the annotated tryptophan metabolites, five were detected in at least 70% of samples. Using linear mixed models across all 711 visits from 237 participants, we tested whether the rate of change in resilience with age differed between resilience groups. Of these, kynurenine showed a significantly steeper age-related increase in the low-resilience group (BH-adjusted *p* = 0.039; **Figure 5a** and **b**). The remaining four metabolites, indole-3-lactate, tryptophan, glucopyranosyl-L-tryptophan, and 5-methylindole-3-carboxaldehyde, showed no significant difference after correction. To examine this at the level of individual participants, we calculated the age slope of kynurenine for each participant separately. Participants with low physical resilience showed steeper slopes than those with high resilience (Wilcoxon rank-sum test, *p* = 0.0062; **Figure 5c**).

**Figure 5.**
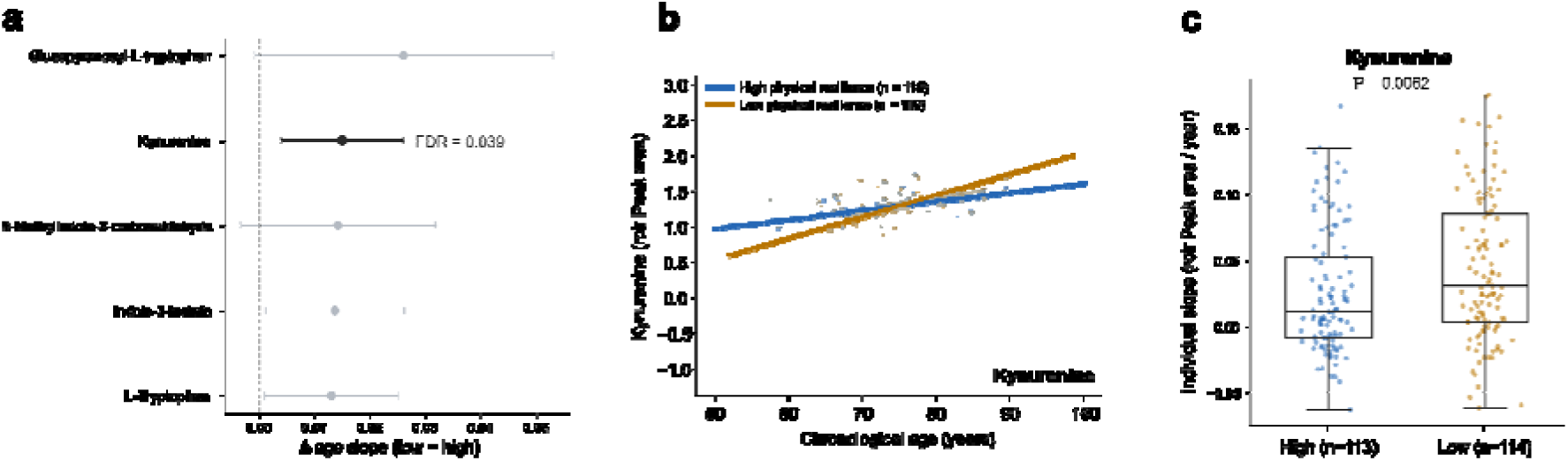
Kynurenine accumulates more steeply with age in individuals with low physical resilience. a) Difference in the age slope between low and high physical resilience groups for tryptophan metabolites detected in at least 70% of samples. Points represent the estimated difference and error bars the 95% confidence interval; metabolites with FDR < 0.05 are shown in black. b) Association of rclr peak area of kynurenine with age (blue = high physical resilience group; yellow = low physical resilience group). All visits of each participant (n=237) are included, and lines represent a linear model fit per group, with the shaded area representing the 95% confidence interval. c) Age slopes calculated for each participant individually, restricted to those with kynurenine detected at all three visits (n=227).

### Dexverapamil Metabolite Mapping Reveals Metabolism-Dependent Associations with Physical Resilience

In the molecular network (**Figure 3a**), several antihypertensive drugs and their metabolites were associated with physical resilience. Dexverapamil, the R-enantiomer of verapamil, was particularly notable, along with its major metabolites norverapamil from N-demethylation and D617 from N-dealkylation. We therefore focused on dexverapamil, a drug whose metabolites were well represented in the dataset, and used MassQL queries to specifically target submolecular networks to systematically capture its metabolite family and examine whether xenobiotic metabolism differs according to physical resilience. Both diagnostic fragment ions arose from cleavage of the C-N bond linking the dimethoxyphenethyl group to the tertiary amine, releasing the nitrogen-containing portion of the molecule as a neutral loss and retaining the dimethoxyphenyl-containing portion as the fragment ion, with *m/z* 150.0675 reflecting an additional loss of a methyl group from *m/z* 165.0910 (**Figure 5a**). Based on these two fragments, we designed a MassQL query to retrieve spectra sharing the dexverapamil substructure across each submolecular network where dexverapamil metabolites were annotated. A second query targeted the glucuronide moiety, a common phase 2 drug metabolic pathway, and identified several dexverapamil metabolites arising from glucuronidation with or without concurrent phase 1 metabolism. For example, a putative metabolite at precursor *m/z* 617.3070 showed both diagnostic verapamil fragments and a neutral loss of 176.0321, indicating that this metabolite arose from glucuronidation combined with demethylation (**Figure 5b**). Applying both queries to the dexverapamil submolecular network revealed a broad set of metabolite nodes linked through combinations of dealkylation, demethylation, hydroxylation, and glucuronidation (**Figure 5c**). Edge color distinguished nodes captured exclusively by the verapamil-targeting query, exclusively by the glucuronide-targeting query, or by both, making it possible to trace overlapping and distinct metabolic modifications within the network. These metabolites were consistently detected together with dexverapamil across individuals (**Figure 5d**). Drug metabolites generally elute earlier than their parent compound due to increased polarity^24^, so we generated extracted ion chromatograms for the dexverapamil metabolites in a representative serum sample to confirm that the network-derived metabolites corresponded to chromatographic features (**Figure 5e**). All annotated metabolites, including those from hydroxylation, demethylation, dealkylation, and combined metabolic modifications, appeared as distinct peaks eluting earlier than dexverapamil, with the demethylation product showing the highest peak intensity. Several isomeric species, such as the dealkylation plus demethylation products eluting at 3.83 and 4.04 min, were baseline-resolved. An *in vitro* metabolism assay using human liver microsomes supplemented with nicotinamide adenine dinucleotide phosphate (NADPH) and uridine 5′-diphosphoglucuronic acid (UDPGA) further confirmed all annotated dexverapamil metabolites, with cosine scores above 0.9 for all matches (**Supplementary Figure S5**). We compared the levels of each metabolite between individuals with high and low physical resilience, normalized to the dexverapamil peak area to account for variability in oral drug and metabolite levels in blood after administration (**Figure 5f**). Metabolites arising from hydroxylation, demethylation, and glucuronidation appeared broadly similar between groups, whereas metabolites involving dealkylation and multiple sequential demethylations tended to be higher in the low resilience group.

### Repository-Scale MS/MS Spectral Searches Uncover Distinct Tissue and Health Status Patterns of Physical Resilience-Associated Features

To further explore where physical resilience-associated features (i.e., 2,810 features selected by the sPLS-R model) are detected across the human body, we examined their distribution across tissue and health status categories using the mass spectrometry search tool (MASST) (**Figure 6**). Depending on the direction of association with physical resilience (i.e., positive or negative), the proportion of associated features detected in each organ differed (**Figure 6a**). We observed positive feature-dominant organs such as heart (93.9%), adipose tissue (73.2%), and liver (71.5%). In contrast, kidney and urine showed markedly lower rates (23.4% and 16.6%). Blood-derived matrices, including plasma and serum, consistently showed a relatively high proportion of positive features (63.5% and 56.4%). Overall, positively associated features tended to be detected more frequently in metabolically active organs, while negatively associated features were more frequently detected in excretory tissues such as the kidney and urine. A potential contrast was also observed across health status categories (**Figure 6b**). These annotations describe the donors of samples deposited in public repositories that are independent of the RBS cohort. While healthy and acute illness samples showed similar proportions of positive and negative features, approximately 60% of features detected in unhealthy (NOS), patient’s disease status is unassigned in ReDU metadata, and chronic illness samples were negatively associated with resilience. Sample sizes differed considerably across groups (healthy samples (n = 1,810) were the most abundant, while acute illness samples (n = 141) were the least represented); however, the consistently higher proportion of negatively associated features in the chronic illness and unhealthy (NOS) groups suggests that these physical resilience-associated features may be more closely associated with sustained illness states than with acute illness.

**Figure 6.**
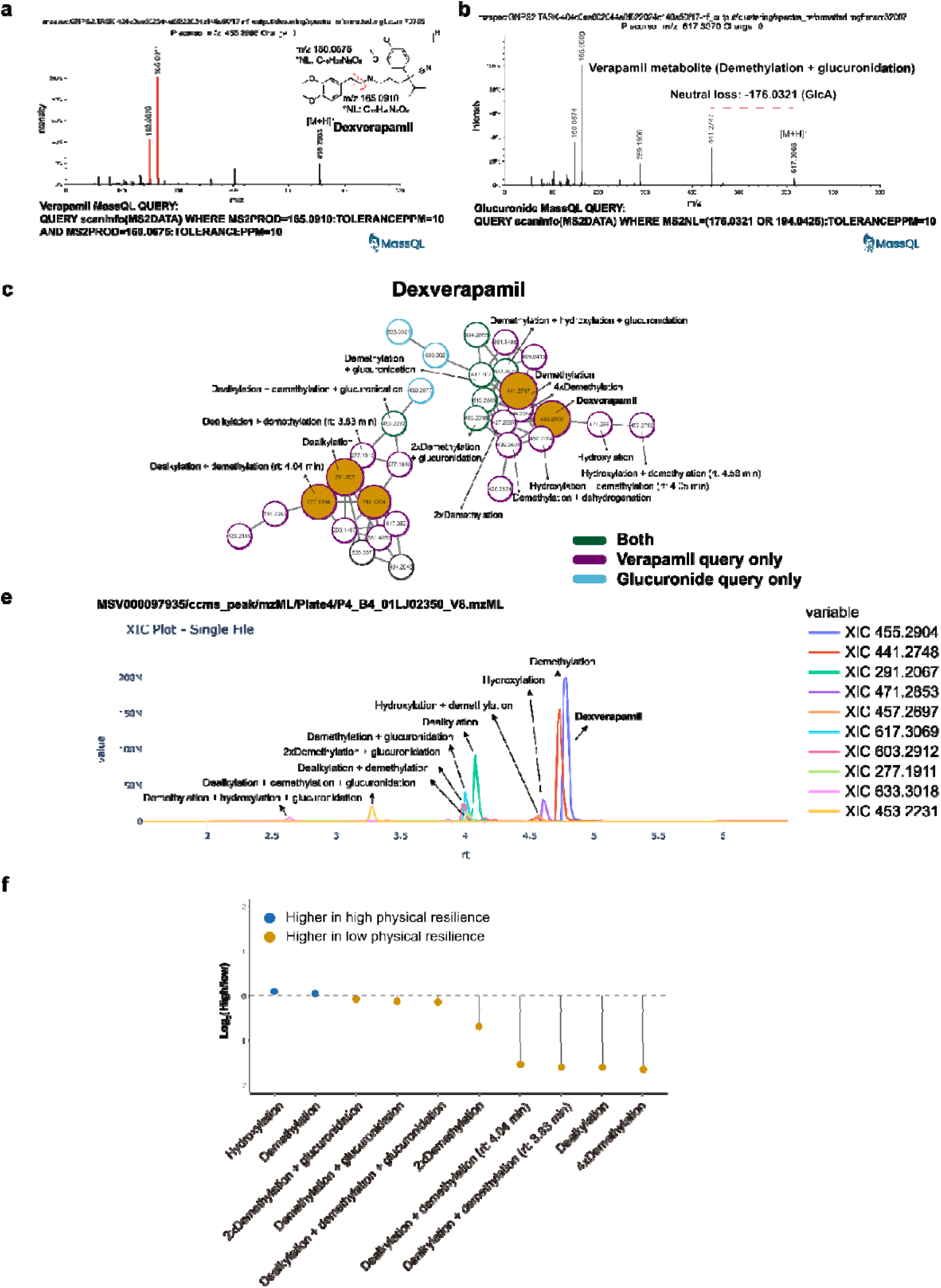
Patterns of dexverapamil metabolites with physical resilience. (a) MS/MS spectra of dexverapamil and the designed MassQL query used to capture its metabolites in the submolecular network. (b) MS/MS spectra of a putative verapamil glucuronide and the designed MassQL query used to target glucuronides. (c) Submolecular network of dexverapamil and its metabolites. Edge colors correspond to the MassQL query results (green: captured by both queries; purple: captured only by the verapamil query; blue: captured only by the glucuronide query). (d) Heatmap showing the abundance of dexverapamil metabolites identified in the submolecular network (only metabolites that co-occurred with dexverapamil are shown). (e) Extracted ion chromatograms of dexverapamil and its metabolites identified in the submolecular network in a serum sample. (f) Bar chart showing the relative abundance of each metabolite across physical resilience groups (values are presented as log2 fold changes, and each metabolite peak area was normalized to the dexverapamil peak area).

## Discussion

In this study, we performed untargeted metabolomic profiling to characterize the molecular signatures of physical resilience in a longitudinal cohort of 237 older adults from the RBS. More than 20,000 metabolic features were detected in serum samples collected across a median of 15.6 years, of which 2,810 were identified by sPLS regression as important contributors to longitudinal variation in physical resilience. Further investigation of these features revealed significant associations across multiple chemical classes, including acylcarnitines, glutamine conjugates, phosphocholines, and several medications. Specifically, acylcarnitines (predominantly medium-chain species), glutamine conjugates, and antihypertensive medications were associated with lower physical resilience, whereas phosphocholines were associated with higher physical resilience (**Figure 3a**). Collectively, these findings identify candidate molecular biomarkers of physical resilience and provide new insights into the biological processes underlying healthy aging. To our knowledge, this is the first study to characterize the molecular signature of physical resilience using an untargeted metabolomics approach. This works contributes to a deeper understanding of the biological mechanisms that support physical resilience may help inform strategies to promote healthy aging and preserve functional independence.

Acylcarnitines are esters of carnitines and acyl groups that play a central role in transporting long-chain fatty acids into mitochondria for β-oxidation and ATP production^25^. Consequently, elevations in circulating acylcarnitines may reflect mitochondrial dysfunction or incomplete β-oxidation. Indeed, previous studies have found higher concentrations of circulating medium- to long-chain acylcarnitines to be predictive of cardiovascular disease^26^, frailty^27^, and increased risk of functional impairment^28^ in older adults. Our findings (**Figure 4a**) are consistent with the literature, demonstrating that primarily medium-chain acylcarnitines were associated with poorer physical resilience, further highlighting the importance of this chemical class in healthy aging. Although the literature remains limited, there is evidence^29^ that regular aerobic exercise may reduce circulating acylcarnitine concentrations by improving mitochondrial functioning and β-oxidation. Future studies should determine whether changes in circulating acylcarnitines mediate improvements in age-related outcomes, but our findings suggest that acylcarnitines may serve as promising biomarkers of physical resilience with potential clinical utility for identifying older adults at risk for physical decline and monitoring physiological responses to exercise interventions. We also found that several glutamine conjugates, particularly fatty acyl- and aromatic acid-conjugated species, were associated with poorer physical resilience (**Figure 4b**). Although the glutamine-conjugated fatty acids (e.g., glutamine C5:0, C6:0, C8:1, C8:2) remain largely understudied, the aromatic acid glutamine conjugates^30^ are well-established products of host-microbial metabolism. Phenylacetylglutamine^30^, one of the most studied metabolites in this pathway, has been associated with numerous cardiovascular, cerebrovascular, and neurological diseases, and may even serve as a predictor of mortality^31^. Our findings extend this literature by demonstrating that glutamine conjugates are also associated with poorer physical resilience in older adults. In contrast, we found that 3 phosphocholines (e.g., L-α-glycerylphosphorylcholine, 1-palmitoylglycerophosphocholine, and a phosphatidylcholine species) were significantly associated with better physical resilience (**Figure 4c**). Phosphocholines are among the most abundant phospholipids in mammalian cell membranes and play essential roles in maintaining membrane integrity, lipid transport, and cellular signaling^32^. Consistent with our findings, a recent systematic review^33^ concluded that reductions in phosphatidylcholine and lysophosphatidylcholine species are characteristic features of aging, suggesting that preservation of phosphocholine metabolism may contribute to healthy aging.

Of the five tryptophan metabolites we examined, only kynurenine differed in its age trajectory between resilience groups, rising more steeply in the low resilience group (**Figure 5**). Kynurenine is produced from tryptophan by indoleamine 2,3-dioxygenase (IDO), an enzyme induced by inflammatory cytokines, and its circulating levels rise with age^34^. In older adults, higher serum kynurenine has been linked to weaker grip strength, slower gait, longer chair-stand times, and a higher risk of frailty, although these findings come from a single time point^23^. Mice fed kynurenine from 16 to 24 months of age, an age range comparable to 56 to 69 years in humans, had reduced endurance capacity and walking speed and a higher prevalence of frailty^22^. Kynurenine also causes bone loss in mice by increasing osteoclast activity and suppressing bone formation^21^. Our data suggest that the rate at which kynurenine accumulates, and not only its level, may differ with physical resilience. Kynurenine levels are also known to respond to lifestyle. Exercise increases skeletal muscle expression of the kynurenine aminotransferases in mice and humans, shifting kynurenine toward kynurenic acid^35^. Diet has a similar effect. In older adults, changes in adherence to a Mediterranean-style diet over 26 weeks were inversely associated with concurrent changes in kynurenine pathway activation^36^, and a methyl-deficient diet raised plasma kynurenine in rats, which choline supplementation partly lowered^37^. Beyond this, the kynurenine signal may also reflect kidney function rather than a musculoskeletal process. Bidirectional Mendelian randomization has shown that reduced estimated glomerular filtration rate (eGFR) raises circulating kynurenine, while kynurenine has no reciprocal effect on eGFR^38^.

In the molecular network, we also noted a cluster of antihypertensive drugs whose parent compounds and metabolites were negatively associated with physical resilience (**Figure 3a**). The presence or absence of these medications in individual subjects did not affect the selected features of the model, indicating that simply taking these drugs was not what drove the resilience-associated metabolite signal (**Supplementary Figure S2**). We examined this further for dexverapamil. Several metabolites from oxidation, including those formed by dealkylation and extensive demethylation, were consistently higher in individuals with low physical resilience, whereas metabolites from hydroxylation, single-step demethylation, and glucuronidation showed little difference between groups (**Figure 6f**). If oxidative metabolism itself were increased in low-resilience individuals, the glucuronide conjugates would also be expected to increase, since they are formed directly from these oxidative metabolites. This was not observed, as glucuronide levels remained largely similar between groups. We therefore reasoned that this pattern is unlikely to reflect a change in drug metabolism capacity itself as a function of physical resilience status, but rather relates to processes occurring after these metabolites are formed. This would be consistent with a decline in renal function, which is known to progress with age and can reduce the elimination of drug metabolites without necessarily altering hepatic metabolic capacity^39^. Although glucuronide conjugates and specific metabolites from one-step oxidation are generally prone to renal excretion, their unaltered levels may reflect differences in renal excretion mechanisms, as such mechanisms are influenced by the physicochemical properties of the metabolites (e.g., glucuronides tend to be excreted via organic anion transporters owing to their greater polarity relative to metabolites from oxidation)^40,41^. A related finding^42^ has been reported for metoprolol in patients with renal impairment, in whom plasma concentrations of the parent drug were unchanged while its metabolite from oxidation accumulated two- to threefold, an effect attributed to reduced renal clearance of the metabolite rather than altered drug metabolism. Hence, our findings may indicate that these differing patterns of drug metabolite abundance, according to physical resilience, could be associated with more impaired renal function in the low physical resilience group, causing accumulation of metabolites from specific types of oxidation. This interpretation parallels our observation for kynurenine, where the steeper age-related increase in the low resilience group may similarly reflect reduced renal clearance.

Notably, the pattern of positively associated features being more frequently detected in highly metabolically active organs such as heart, liver, and adipose tissue may suggest that these features are preferentially associated with tissues engaged in active substrate turnover and energy metabolism (**Figure 7a**). This raises the possibility that maintenance of active metabolic function in these organs is one of the physiological correlates of higher physical resilience^43–45^. Negatively associated features being more frequently detected in organs responsible for excretion^46^ (**Figure 7a**), such as the kidney and urine, may suggest that these features are preferentially associated with processes involved in the clearance of metabolic byproducts that could be correlated with lower physical resilience^47^.

**Figure 7.**
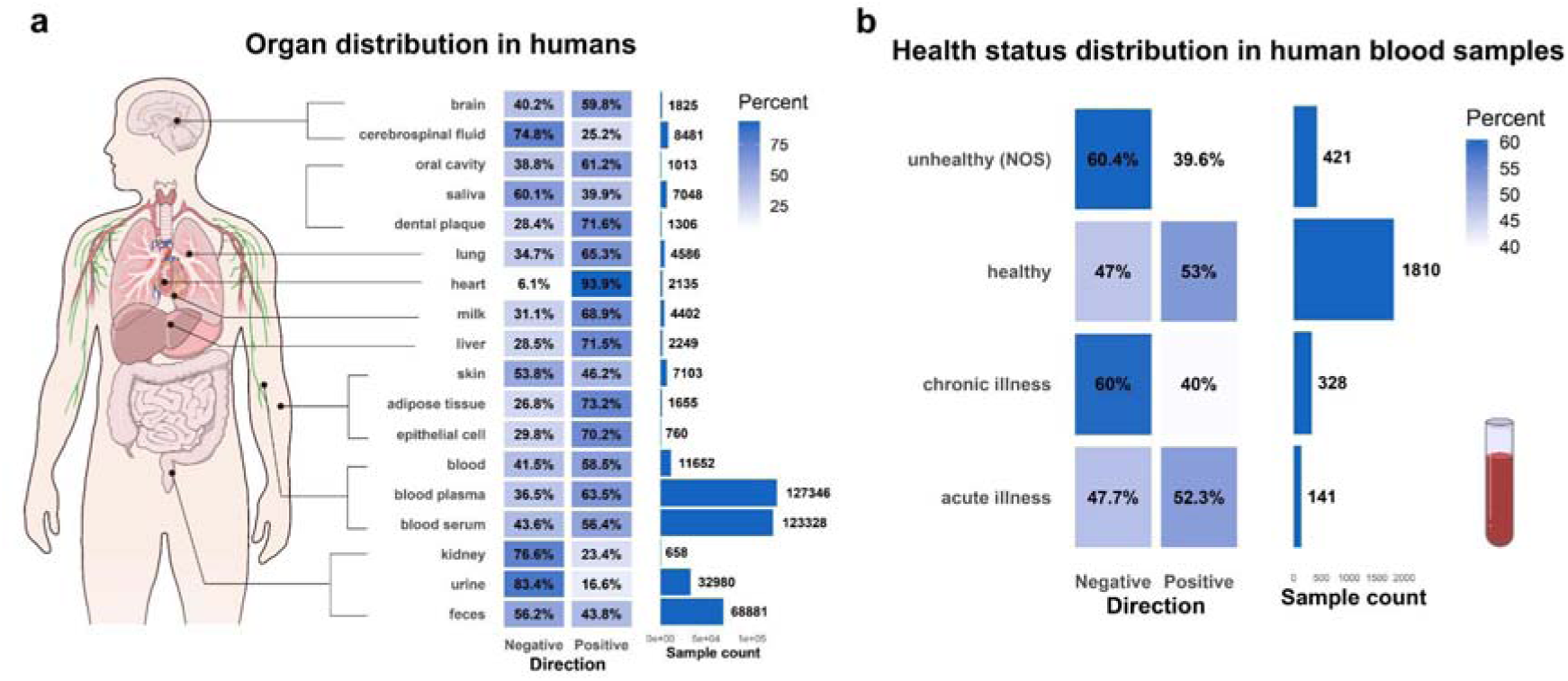
Repository-scale mapping of physical resilience-associated features across tissues and health status using MASST. (a) Distribution of matched features across human tissue matrices. (b) Distribution of matched features across human health status categories. NOS, not otherwise specified (used when a specific disease status is not assigned to the patient in the ReDU metadata).

## Conclusion

This study provides the first untargeted metabolomic characterization of physical resilience in a community-based longitudinal cohort, identifying acylcarnitines, glutamine conjugates, phosphocholines, and tryptophan metabolites as candidate molecular correlates of physical resilience status. Kynurenine showed a steeper age-related increase in individuals with low physical resilience, and metabolites of the antihypertensive drug dexverapamil arising from oxidation, but not from glucuronidation, tended to be higher in this group. These associations point to mitochondrial fatty acid oxidation, membrane lipid homeostasis, and possibly elimination capacity as processes relevant to physical resilience. Repository-scale search further supports that the physical resilience-associated features show varying detection frequency across distinct tissues, highlighting the involvement of both actively metabolizing organs and excretory tissues in these signatures. Together, these findings offer testable biomarker candidates and mechanistic hypotheses that may inform future strategies to monitor and support physical resilience across the aging process.

### Limitations of the study

This study has several limitations. First, the RBS is a relatively homogeneous cohort, and our findings require replication in independent and more diverse populations. Second, although drug metabolites were confirmed by *in vitro* enzymatic reactions, a large portion of the metabolites reported from this untargeted workflow were putatively annotated by spectral library matching (level 2-3 according to Metabolomics Standard Initiative) or *in silico* prediction, and confirmation with authentic standards will be needed in follow-up studies. Third, the drug metabolism findings were derived from individuals in whom the respective drugs were detected and should be considered exploratory. Fourth, kidney function was not measured in this cohort, and the interpretation that reduced renal clearance contributes to the observed patterns for kynurenine and dexverapamil metabolites remains untested. Lastly, MASST results only reflect a spectral match in public repositories rather than a feature’s tissue of origin or causal role. Thus, detection in metabolically active organs or excretory tissues should not be interpreted as directly indicating benefit or harm to physical resilience.

## Materials availability

This study did not generate unique reagents.

## Lead contact

Further information and requests for resources and reagents should be directed to the lead contact, Anthony Molina.

## Data and code availability

The untargeted LC-MS/MS data is publicly available on GNPS/Massive under the accession MSV000097935. The corresponding GNPS2 job is available at https://gnps2.org/status?task=404c0ee602644a6f822024c146a50f17. Code and scripts used for analysis in this manuscript is available on GitHub: https://github.com/jeonginseo95/Aging_physical_resilience.

## Acknowledgements

This work was funded by The Wellcome Leap Dynamic Resilience program (co-funded by Temasek Trust). Data collection for the Rancho Bernardo Study of Healthy Aging was provided primarily by the National Institutes of Health (including grant numbers: HV012160, AA021187, AG028507, AG007181, DK31801, HL034591, HS06726 and HL089622). Archiving and sharing of RBS data was supported by the National Institute on Aging: AG054067. J.I.S. is supported by the National Research Foundation of Korea (NRF) (RS-2025-02373133). T.S. is supported by the Fulbright Visiting Scholar Program through the Fulbright NL Promovendus Grant, which is sponsored by the U.S. Department of State and the Fulbright Netherlands Commission.

## Author contributions

Conceptualization, I.M., P.C.D., A.M.; methodology, P.C.D., I.M., J.I.S., T.S., J.B; formal analysis, I.M., J.I.S., T.S.; investigation, I.M., J.I.S., T.S., C.W., K.K., W.N., J.Z., J.B.; resources, J.B., A.M.; writing -original draft, I.M., P.C.D., A.M., J.I.S., T.S.; writing -review & editing, all authors; supervision, I.M., P.C.D., funding acquisition, P.C.D., A.M.

## Methods

### Study population

Samples were collected from the Rancho Bernardo Study of Healthy Aging^48^, a prospective population based cohort study initiated in 1972, in Rancho Bernardo. People enrolled were minimally 30 years old and included between 1972 and 1974. 6,726 residents (82% of the adult population in Rancho Bernardo)^49^ were included and followed up at 4-year intervals for >40 years^49^. We selected a subset of 237 participants for metabolomics analysis based on availability of 3 serum samples and physical resilience scores. There were no exclusion criteria. All study visits were approved by the University of California San Diego Human Research Protections Program (UCSD IRB 960019; UCSD IRB 040165; UCSD IRB 130757). All participants provided written informed consent at each clinic visit.

### Calculation of physical resilience score

These physical resilience scores were calculated based on a participant’s performance on the Grip Strength test and the Timed Chair stand^14^. 2,630 participants were included in the calculation of physical performance trajectories. A linear mixed effects-regression model was built on the composite z-score of these tests for these participants, using fixed effects (age, sex and education) and random participant-level intercepts and slopes. The overall model slope was -0.49, indicative of a 0.49 SD decrease in composite physical function score per decade after 70 years of age.

### Sample collection and preparation

Blood samples were obtained via venipuncture after a requested 12-hour fast. Serum was separated and frozen at -70 in sunlight-protected tubes. Serum samples were processed using the Phenomenex Phree™ 96-well phospholipid removal kit (Phenomenex Inc., Torrance, CA, USA), maintained at 4:1 ration using 100% methanol, and dried and concentrated using a CentriVap Benchtop Vacuum Concentrator (Labconco, Kansas City, MO, USA). as per previous protocol^11^.

### LC-MS/MS data acquisition and data processing

LC-MS/MS methods are described in previous work^11,50^. Briefly, analysis was performed on Thermo Scientific™ Q Exactive™ Mass Spectrometer system coupled with a Thermo Scientific™ Vanquish™ UHPLC System (Thermo Scientific, Waltham, MA). Chromatographic separation was performed on a Phenomenex Luna Omega Polar C18 HPLC column (2.1 x 50 mm, 1.6 μm; Torrence, CA). After acquiring the MS/MS spectra, they were converted to mzML files using MSconvert (ProteoWizard)^51^ and deposited in GNPS/MassIVE under: MSV000097935. Feature extraction was done using MZmine 4.20. The feature list was then exported as a feature quantification table (.csv) and an MGF spectra file. Metabolite annotations were performed with the default GNPS Spectral Libraries and the GNPS-BILE-ACID-MODIFICATIONS, GNPS-MASSQL-BILE-ACID-ISOMER, Carnitines_library_2025_testing_V2_MassQL_only and the MULTIPLEX-SYNTHESIS-FILTERED libraries using the Feature-Based Molecular Networking (FBMN) workflow on GNPS2 and the job can be found at: https://gnps2.org/status?task=404c0ee602644a6f822024c146a50f17. The network.graphml file from FBMN was imported into Cytoscape (version 3.10.3)^52^ for visualization and analysis. MS/MS spectra of features without library annotation were imported into SIRIUS (version 6.3.2)^53^, and CANOPUS was used to predict their chemical classes; predicted classes were retained only if the probability score was > 0.7. Individual features that were statistically significant within each chemical family shown in **Figure 4**, as well as all features selected by the sPLS-R model (**Figure 6**), were searched against public repositories using the MASST^54^ tool (i.e., domainMASST) within the GNPS ecosystem, with their MS/MS spectra used as input, to assess their relevance to specific diseases and their distribution across organs. The following parameters were used to define a match: a minimum of 4 matching peaks and cosine similarity > 0.7, with precursor and fragment ion mass tolerances set at 0.02 Da.

### Sensitivity analysis for sex and BMI

To assess whether the unequal sex distribution between resilience groups influenced the results, features differing between resilience groups were identified by Wilcoxon rank-sum tests after rclr transformation, and the comparison was repeated separately within each sex and using a Wilcoxon rank-sum test stratified on sex, all corrected for multiple testing using Benjamini-Hochberg false discovery rate. Effect sizes were quantified using Cliff’s δ and compared between sexes by Pearson correlation. BMI was compared between resilience groups within each sex using a Wilcoxon rank-sum test.

### sPLS-R model development

Data preprocessing was performed as described in earlier work^11^. Briefly, feature and annotation data was imported in R 4.4.3. Features eluting before 0.7 minutes were excluded, sample quality was assessed using inspection of total intensity profiles, and we subtracted features that did not reach 5 times the peak area of blanks samples. Finally, we applied RCLR conversion using package vegan 2.6-10, and removed near-zero variance features. We built a parse partial least squares regression (sPLS-R) model using package mixOmics 6.30. The model was tuned using mixOmics 6.30 and inflection 1.3.7. We evaluated explained variance with linear modeling, performance with 10-fold cross validation and stability with permutation testing (n = 100).

### Statistical analysis

For univariate analysis, the association between features and physical resilience were assessed using Wilcoxon tests. Associations of features with age were assessed using linear mixed modeling, performed using package lme4 (version 1.1-37) and lmerTest (version 3.1-3). The carnitines and phosphocholines of interest were selected for non-sparsity, cut-off at <70%. Zero-values were handled using a feature-specific pseudocount, at half the minimum non-zero intensity. Finally, data was log2 transformed. Centered age was used as a predictor, with participants as random intercept and random slope. Significance for linear mixed modeling and univariate analysis was corrected for multiple testing using Benjamini-Hochberg false discovery rate.

### *In vitro* enzymatic reaction

The *in vitro* enzymatic reaction followed previously reported procedures^55,56^. The reaction system consisted of human liver microsomes pooled from 50 donors (1 mg/mL; Gibco™, Grand Island, NY), 0.1 M potassium phosphate buffer (pH 7.4; Fisher Scientific, Pittsburgh, PA), 2.5 mM MgCl2 (Alfa Aesar, Ward Hill, MA), and 25 μg/mL alamethicin (Cayman Chemical, Ann Arbor, MI). Verapamil (100 μM; Combi-Blocks, San Diego, CA) dissolved in DMSO as a stock solution was incubated in this system. After pre-incubation at 37 °C for 5 min, the reaction was initiated by adding NADPH generating system (10 mg/mL NADP (ChemCruz, Dallas, TX)), 0.1 M glucose-6-phosphate (Thermo Scientific, Waltham, MA), and 0.1 M of glucose-6-phosphate dehydrogenase (Sigma-Aldrich, St. Louis, Missouri)) and uridine 5-diphosphoglucuronic acid (2 mM) into the reaction mixture. The total reaction volume was maintained at 200 μL. After 2 h of incubation, the reaction was terminated by adding 100 μL of ice-cold methanol. The samples were then vortexed for 1 min, centrifuged at 13,200 rpm for 5 min, and the resulting supernatants were subjected to LC-MS/MS analysis.

## Disclosures

PCD is an advisor and holds equity in Cybele, BileOmix and Sirenas and a Scientific co-founder, advisor and holds equity to Ometa, Enveda, and Arome with prior approval by UC-San Diego. PCD also consulted for DSM animal health in 2023. In the Dorrestein Lab, the use of AI, generative AI, and large language models (LLMs), and software that uses these technologies, both free and commercial, is encouraged across all aspects of research, including literature review, coding, data analysis, and text editing. All figures and analyses are original. For transparency and reproducibility, all raw data, derived data tables and final code used in this research are made accessible and linked with this manuscript.

## References

1. Hoogendijk, E. O. et al. Frailty: implications for clinical practice and public health. The Lancet 394, 1365–1375 (2019).

2. Mathers, C. D., Stevens, G. A., Boerma, T., White, R. A. & Tobias, M. I. Causes of international increases in older age life expectancy. The Lancet 385, 540–548 (2015).

3. Ferrucci, L. & Kuchel, G. A. Heterogeneity of Aging: Individual Risk Factors, Mechanisms, Patient Priorities, and Outcomes. J. Am. Geriatr. Soc. 69, 610–612 (2021).

4. Whitson, H. E. et al. Physical Resilience in Older Adults: Systematic Review and Development of an Emerging Construct. J. Gerontol. Ser. A 71, 489–495 (2016).

5. Resnick, B., Galik, E., Dorsey, S., Scheve, A. & Gutkin, S. Reliability and Validity Testing of the Physical Resilience Measure. The Gerontologist 51, 643–652 (2011).

6. Zhang, H. et al. Assessment of Physical Resilience Using Residual Methods and Its Association With Adverse Outcomes in Older Adults. Innov. Aging 7, igad118 (2023).

7. Stark, L. et al. Physical Resilience May Offset Mortality Risks Associated With Genetic Predisposition to Shorter Survival: A Population-based Cohort Study. J. Gerontol. Ser. A 80, glaf101 (2025).

8. Panyard, D. J., Yu, B. & Snyder, M. P. The metabolomics of human aging: Advances, challenges, and opportunities. Sci. Adv. 8, eadd6155 (2022).

9. Ramírez-Vélez, R. et al. Lipidomic signatures from physically frail and robust older adults at hospital admission. GeroScience 44, 1677–1688 (2022).

10. McEvoy, L. et al. THE RANCHO BERNARDO STUDY (RBS) OF HEALTHY AGING: A RICH RESOURCE FOR STUDYING AGING IN WOMEN. Innov. Aging 3, S355 (2019).

11. Scheurink, T. A. W. et al. Serum metabolic signatures of cognitive resilience in a longitudinal aging cohort. 2026.03.29.715122 Preprint at 10.64898/2026.03.29.715122 (2026).

12. Jarmusch, A. K. et al. ReDU: a framework to find and reanalyze public mass spectrometry data. Nat. Methods 17, 901–904 (2020).

13. El Abiead, Y., et al. Structure-centric searching enables global mapping of the public metabolome. Nat. Biotechnol. 1–6 (2026) doi:10.1038/s41587-026-03082-8.

14. Rossiter-Fornoff, J. E., Wolf, S. L., Wolfson, L. I., Buchner, D. M., & FICSIT Group. A Cross-sectional Validation Study of the FICSIT Common Data Base Static Balance Measures. J. Gerontol. Ser. A 50A, M291–M297 (1995).

15. Katzman, W. B., Huang, M.-H., Kritz-Silverstein, D., Barrett-Connor, E. & Kado, D. M. Diffuse Idiopathic Skeletal Hyperostosis (DISH) and Impaired Physical Function: The Rancho Bernardo Study. J. Am. Geriatr. Soc. 65, 1476–1481 (2017).

16. Nothias, L.-F. et al. Feature-based molecular networking in the GNPS analysis environment. Nat. Methods 17, 905–908 (2020).

17. Aron, A. T. et al. Reproducible molecular networking of untargeted mass spectrometry data using GNPS. Nat. Protoc. 15, 1954–1991 (2020).

18. Dambrova, M., et al. Acylcarnitines: Nomenclature, Biomarkers, Therapeutic Potential, Drug Targets, and Clinical Trials. Pharmacol. Rev. 74, 506–551 (2022).

19. Dührkop, K. et al. SIRIUS 4: a rapid tool for turning tandem mass spectra into metabolite structure information. Nat. Methods 16, 299–302 (2019).

20. Zuffa, S. et al. A multi-organ metabolomics atlas reveals molecular dysregulations in Alzheimer’s disease mouse models. Cell Rep. 45, (2026).

21. Refaey, M. E. et al. Kynurenine, a Tryptophan Metabolite That Accumulates With Age, Induces Bone Loss. J. Bone Miner. Res. 32, 2182–2193 (2017).

22. Kawaida, M. Y., et al. Elevating Circulating L-Kynurenine Promotes Frailty in Aging Mice. J. Cachexia Sarcopenia Muscle 17, e70214 (2026).

23. Jang, I.-Y. et al. The association of circulating kynurenine, a tryptophan metabolite, with frailty in older adults. Aging 12, 22253–22265 (2020).

24. Jiang, F. et al. Signal interference between drugs and metabolites in LC-ESI-MS quantitative analysis and its evaluation strategy. J. Pharm. Anal. 14, 100954 (2024).

25. McCoin, C. S., Knotts, T. A. & Adams, S. H. Acylcarnitines—old actors auditioning for new roles in metabolic physiology. Nat. Rev. Endocrinol. 11, 617–625 (2015).

26. Guasch-Ferré, M. et al. Plasma acylcarnitines and risk of cardiovascular disease: effect of Mediterranean diet interventions1, 2, 2. Am. J. Clin. Nutr. 103, 1408–1416 (2016).

27. Meng, L. et al. Specific Metabolites Involved in Antioxidation and Mitochondrial Function Are Correlated With Frailty in Elderly Men. Front. Med. 9, (2022).

28. Caballero, F. F. et al. Plasma acylcarnitines and risk of lower-extremity functional impairment in older adults: a nested case–control study. Sci. Rep. 11, 3350 (2021).

29. Rodríguez-Gutiérrez, R. et al. Impact of an exercise program on acylcarnitines in obesity: a prospective controlled study. J. Int. Soc. Sports Nutr. 9, 22 (2012).

30. Krishnamoorthy, N. K. et al. Role of the Gut Bacteria-Derived Metabolite Phenylacetylglutamine in Health and Diseases. ACS Omega 9, 3164–3172 (2024).

31. Deelen, J. et al. A metabolic profile of all-cause mortality risk identified in an observational study of 44,168 individuals. Nat. Commun. 10, 3346 (2019).

32. McMaster, C. R. From yeast to humans –roles of the Kennedy pathway for phosphatidylcholine synthesis. FEBS Lett. 592, 1256–1272 (2018).

33. Zarezadeh, M. et al. Serum phospholipids during aging: A comprehensive systematic review of cross-sectional and case-control studies. Health Promot. Perspect. 15, 23–36 (2025).

34. Bakker, L. et al. Relation of the kynurenine pathway with normal age: A systematic review. Mech. Ageing Dev. 217, 111890 (2024).

35. Agudelo, L. Z. et al. Skeletal Muscle PGC-1α1 Modulates Kynurenine Metabolism and Mediates Resilience to Stress-Induced Depression. Cell 159, 33–45 (2014).

36. Beers, S. et al. Association Between the Dutch Mediterranean-Dietary Approaches to Stop Hypertension Intervention for Neurodegenerative Delay (MIND-NL) Diet Adherence and Systemic Tryptophan Metabolites in Older Adults at Risk of Cognitive Decline: An Exploratory Study. Mol. Nutr. Food Res. 70, e70377 (2026).

37. Tillmann, S. et al. The Kynurenine Pathway Is Upregulated by Methyl-deficient Diet and Changes Are Averted by Probiotics. Mol. Nutr. Food Res. 65, 2100078 (2021).

38. Cheng, Y. et al. The relationship between blood metabolites of the tryptophan pathway and kidney function: a bidirectional Mendelian randomization analysis. Sci. Rep. 10, 12675 (2020).

39. Musso, C. G. & Oreopoulos, D. G. Aging and Physiological Changes of the Kidneys Including Changes in Glomerular Filtration Rate. Nephron Physiol. 119, p1–p5 (2011).

40. Järvinen, E. et al. The Role of Uptake and Efflux Transporters in the Disposition of Glucuronide and Sulfate Conjugates. Front. Pharmacol. 12, 802539 (2022).

41. Ito, S. et al. Relationship Between the Urinary Excretion Mechanisms of Drugs and Their Physicochemical Properties. J. Pharm. Sci. 102, 3294–3301 (2013).

42. Lloyd, P., John, V. A., Signy, M. & Smith, S. E. The effect of impaired renal function on the pharmacokinetics of metoprolol after single administration of a 1419014190 metoprolol OROS system. Am. Heart J. 120, 478–482 (1990).

43. Kim, J. B. Dynamic cross talk between metabolic organs in obesity and metabolic diseases. Exp. Mol. Med. 48, e214–e214 (2016).

44. Luo, L. & Liu, M. Adipose tissue in control of metabolism. J. Endocrinol. 231, R77–R99 (2016).

45. Mietus-Snyder, M., et al. Next Generation, Modifiable Cardiometabolic Biomarkers: Mitochondrial Adaptation and Metabolic Resilience: A Scientific Statement From the American Heart Association. Circulation 148, 1827–1845 (2023).

46. Schlosser, P. et al. Genetic studies of urinary metabolites illuminate mechanisms of detoxification and excretion in humans. Nat. Genet. 52, 167–176 (2020).

47. Chao, C.-T. & Lin, S.-H. Uremic Toxins and Frailty in Patients with Chronic Kidney Disease: A Molecular Insight. Int. J. Mol. Sci. 22, 6270 (2021).

48. Barrett-Connor, E. Why Women Have Less Heart Disease Than Men and How Diabetes Modifies Women’s Usual Cardiac Protection: A 40-Year Rancho Bernardo Cohort Study. Glob. Heart 8, (2013).

49. McEvoy, L. et al. THE RANCHO BERNARDO STUDY (RBS) OF HEALTHY AGING: A RICH RESOURCE FOR STUDYING AGING IN WOMEN. Innov. Aging 3, S355 (2019).

50. Mohanty, I. et al. The underappreciated diversity of bile acid modifications. Cell 187, 1801–1818.e20 (2024).

51. Chambers, M. C. et al. A cross-platform toolkit for mass spectrometry and proteomics. Nat. Biotechnol. 30, 918–920 (2012).

52. Shannon, P. et al. Cytoscape: A Software Environment for Integrated Models of Biomolecular Interaction Networks. Genome Res. 13, 2498–2504 (2003).

53. Dührkop, K. et al. SIRIUS 4: a rapid tool for turning tandem mass spectra into metabolite structure information. Nat. Methods 16, 299–302 (2019).

54. Wang, M. et al. Mass spectrometry searches using MASST. Nat. Biotechnol. 38, 23–26 (2020).

55. Ha, E. J. et al. Preclinical Bioavailability Assessment of a Poorly Water-Soluble Drug, HGR4113, Using a Stable Isotope Tracer. Pharmaceutics 15, 1684 (2023).

56. Seo, J. I., Jin, G. & Yoo, H. H. Pharmacokinetic considerations for enhancing drug repurposing opportunities of anthelmintics: Niclosamide as a case study. Biomed. Pharmacother. 173, 116394 (2024).

