## Supplementary Figures S1-S5 for "Serum metabolomics reveals signatures associated with physical resilience trajectories from middle to older age"

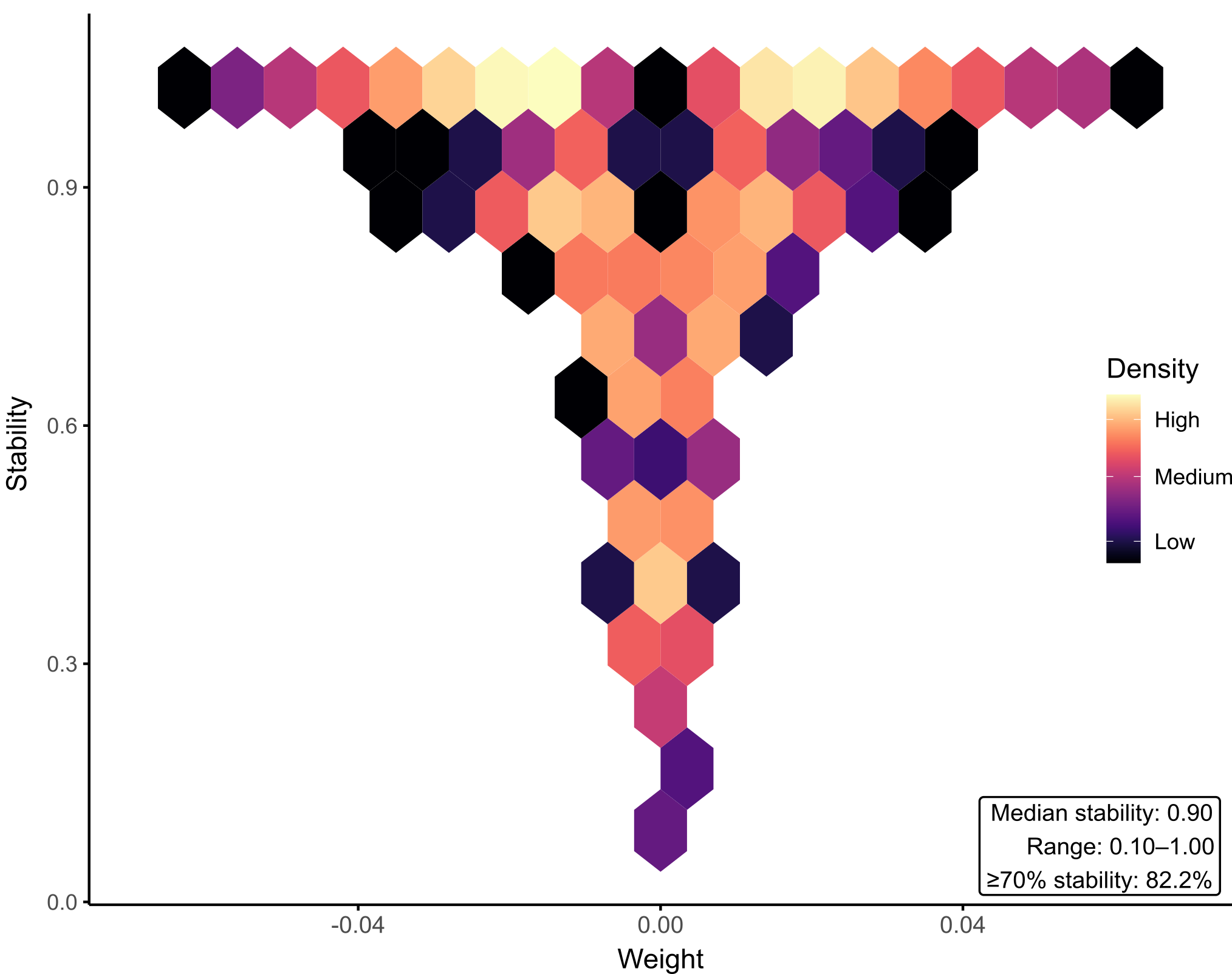

**Supplementary Figure S1.** Stability of selected features across 10-fold cross-validation. A hexplot visualizing the density of features across sPLS weight and selection stability. The inset box summarizes stability metrics.

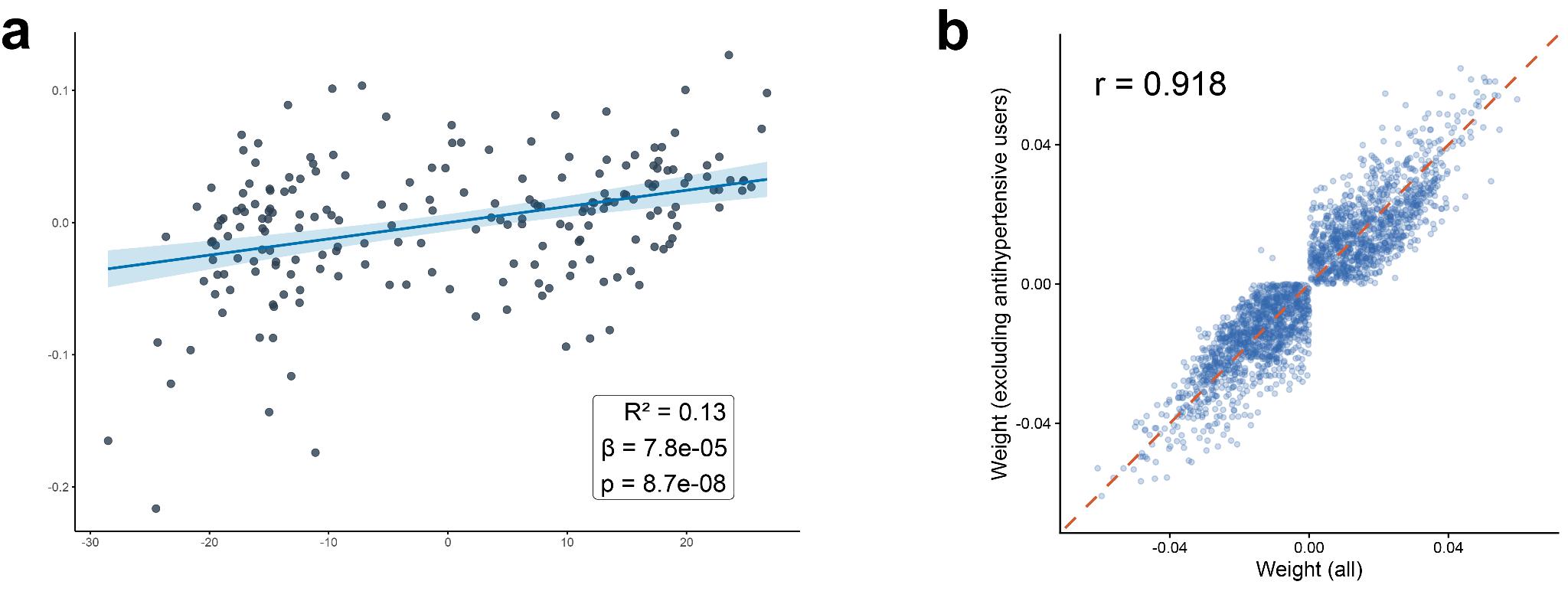

**Supplementary Figure S2.** **Analysis excluding individuals in whom antihypertensive medications were detected, to assess the robustness of the association between metabolite profile and physical resilience.** (a) A linear regression of the first component of the sPLS model against the physical resilience scores. (b) A Pearson correlation plot evaluating the similarity of metabolite profiles between analyses including all individuals (n = 237) and excluding individuals using antihypertensive medications (n = 202).

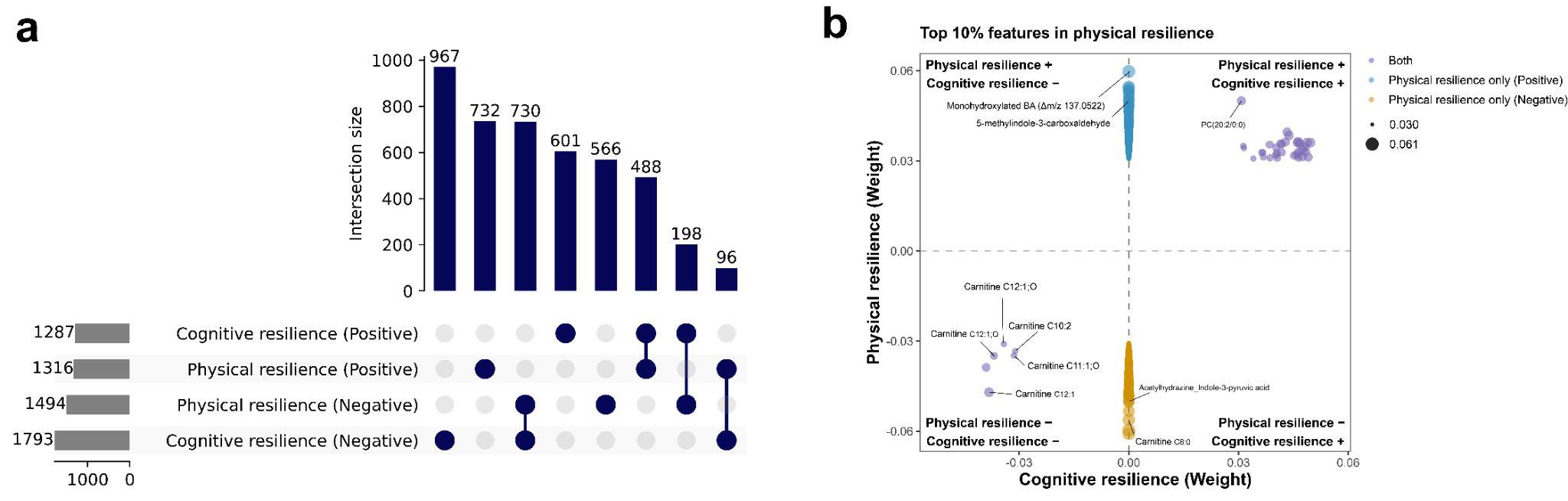

**Supplementary Figure S3. Shared and distinct metabolite features associated with physical and cognitive resilience.** (a) UpSet plot showing the overlap of metabolite features positively and negatively associated with physical and cognitive resilience^1^. (b) Quadrant plot showing the distribution of the top 10% metabolite features selected by the physical and cognitive resilience sPLS models. Feature weights from the cognitive and physical resilience models are plotted on the x- and y-axes, respectively. Node size corresponds to feature weight, and GNPS2 library annotations are shown where available.

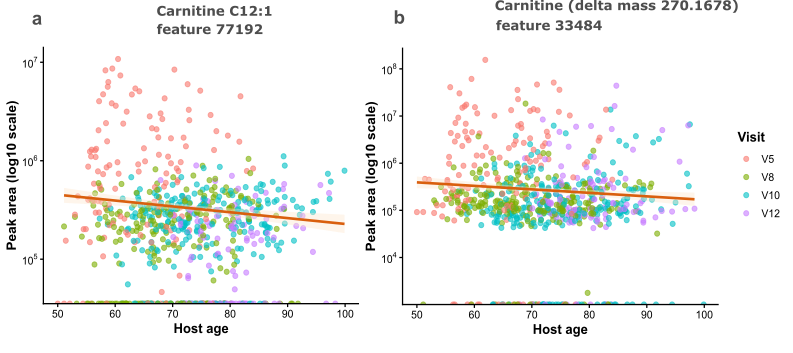

**Supplementary Figure S4. Distribution of abundant carnitines across age and study visits.** Association of log transformed peak area of a) carnitine C12:1 and b) unannotated carnitine (delta mass 270.1678) with age. All 3 visits of each participant are included, and the dots are colored per visit. The line represents a linear model with the shaded area representing the 95% confidence interval.

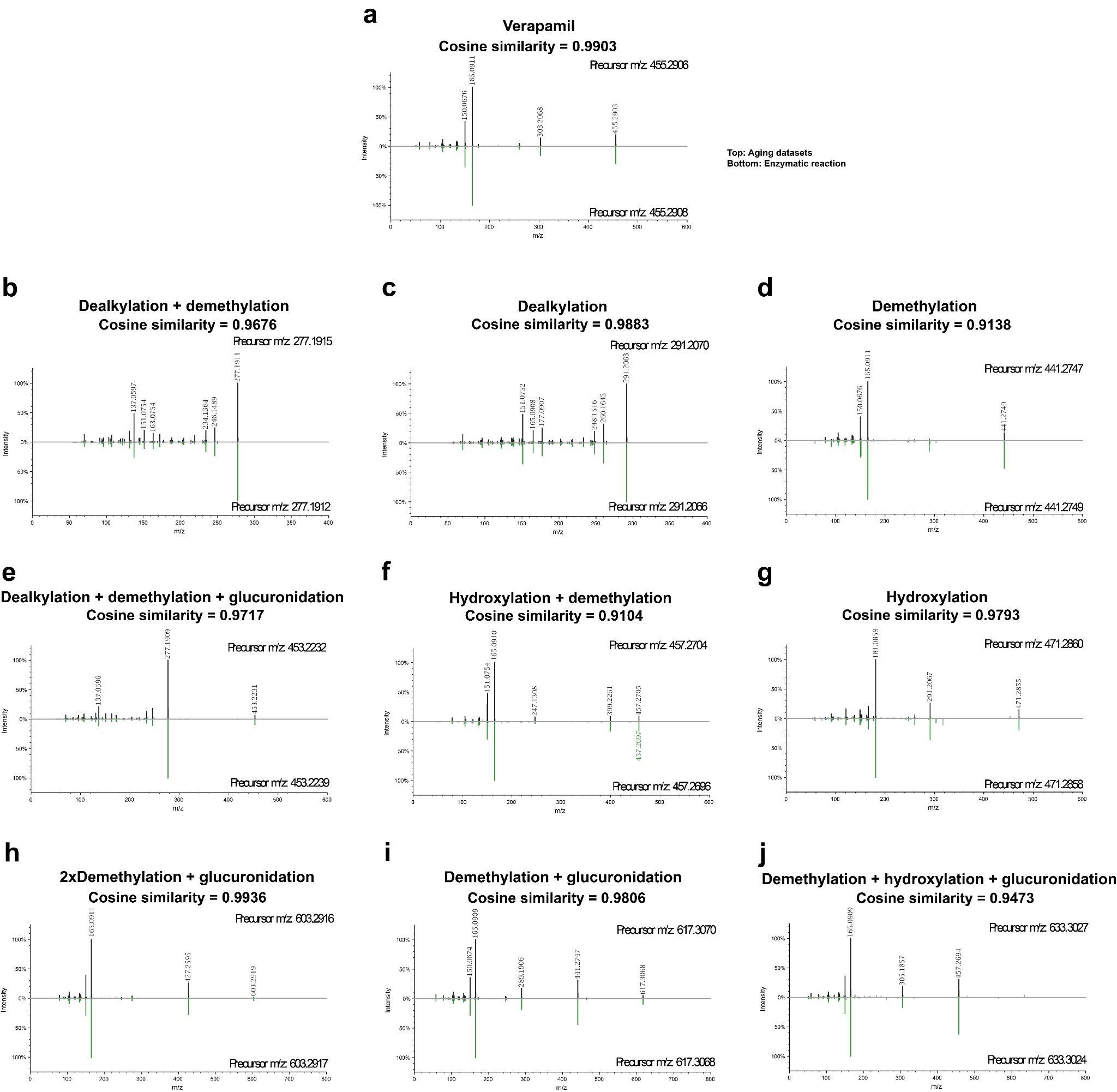

**Supplementary Figure S5.** Mirror plots of (Dex)verapamil and its putative metabolites identified in the RBS cohort. MS/MS spectral similarities between metabolites detected in the RBS cohort (top spectra) and enzymatically generated metabolites (bottom spectra) are shown for (a) (Dex)verapamil and metabolites potentially formed through (b) dealkylation + demethylation, (c) dealkylation, (d) demethylation, (e) dealkylation + demethylation + glucuronidation, (f) hydroxylation + demethylation, (g) hydroxylation, (h) 2×demethylation + glucuronidation, (i) demethylation + glucuronidation, and (j) demethylation + hydroxylation + glucuronidation.

**References**

1.	Scheurink, T. A. W. *et al.* Serum metabolic signatures of cognitive resilience in a longitudinal aging cohort. 2026.03.29.715122 Preprint at https://doi.org/10.64898/2026.03.29.715122 (2026).
