## Supplementary Table 1 for "Serum metabolomics reveals signatures associated with physical resilience trajectories from middle to older age"

**Supplementary Table 1.** Demographics of the selected cohort per resilience group.

| **Characteristic** | **Overall**   N = 237^1^ | **High resilience**   N = 118^1^ | **Low resilience**   N = 119^1^ | **p**^2^ |
| --- | --- | --- | --- | --- |
| **Age** | 64.29 (7.20) | 63.74 (7.66) | 64.84 (6.70) | 0.12 |
| **Sex (Female)** | 152 (64%) | 90 (76%) | 62 (52%) | <0.001 |
| **BMI** | 25.34 (3.47) | 24.82 (3.35) | 25.86 (3.53) | 0.012 |
| BMI – Males | 26.74 (2.96) | 26.15 (2.69) | 27.03 (3.07) | 0.2 |
| BMI – Females | 24.56 (3.50) | 24.41 (3.44) | 24.78 (3.62) | 0.4 |
| **Physical resilience score** | 0.00 (0.05) | 0.03 (0.03) | -0.04 (0.04) | <0.001 |
| ^1^Mean (SD); n (%). ^2^Wilcoxon rank sum test; Pearson's Chi-squared test. BMI = body mass index. | | | | |
